# Unsupervised deep learning with variational autoencoders applied to breast tumor genome-wide DNA methylation data with biologic feature extraction

**DOI:** 10.1101/433763

**Authors:** Alexander J. Titus, Owen M. Wilkins, Carly A. Bobak, Brock C. Christensen

## Abstract

Recent advances in deep learning, particularly unsupervised approaches, have shown promise for furthering our biological knowledge through their application to gene expression datasets, though applications to epigenomic data are lacking. Here, we employ an unsupervised deep learning framework with variational autoencoders (VAEs) to learn latent representations of the DNA methylation landscape from three independent breast tumor datasets. Through interrogation of methylation-based learned latent dimension activation values, we demonstrate the feasibility of VAEs to track representative differential methylation patterns among clinical subtypes of tumors. CpGs whose methylation was most correlated VAE latent dimension activation values were significantly enriched for CpG sparse regulatory regions of the genome including enhancer regions. In addition, through comparison with LASSO, we show the utility of the VAE approach for revealing novel information about CpG DNA methylation patterns in breast cancer.

## Introduction

Deep learning has revolutionized computer vision and natural language and speech processing but has seen relatively slow adoption to biological datasets [1]. This slow adoption is due, in part, to a deficit of well annotated datasets of a sufficient size. Furthermore, extraction of specific biological features from learned models remains challenging. In the case of building supervised learning algorithms capable of predicting biological or phenotypic states, models depend on training data with labels such as “case-control” status or “tumor status” based on an existing cohort. However, to investigate the utility of learning algorithms for discovery-based approaches on biological data features where validated genomic annotations are not yet defined, supervised methods that broadly categorize data are not suitable. Instead, unsupervised algorithms (e.g. unsupervised hierarchical clustering), are promising alternatives, and recently, unsupervised deep learning models have started to emerge. However, new approaches are required to characterize the biology captured in outputs from unsupervised deep learning models. Efficient extraction of relevant biological information from learned features is critical for determination of whether new unsupervised learning methods are poised to address limitations of other approaches to analyze genome-scale molecular data sets.

Variational autoencoders (VAE) are an unsupervised deep learning approach that has garnered extensive recent interest. While supervised models learn by testing predicted labels against so-called ‘ground truth’ labels, for example “case vs control”, VAEs learn through data reconstruction [2]. Specifically, VAEs take raw data as input, learn latent representations and distributions of the input data, and then sample from the learned distribution to reconstruct the original input data. Efficacy of the learned model is then determined through comparing the reconstructed data to the original input. VAEs combine input data and data between hidden layers in a way that can be thought of as a high dimensional interaction space. Each of the input data features are combined in weighted proportions that contribute to the learned latent dimensions of the models. This in turn provides an opportunity to study genome-wide interactions in ways that may be computationally intractable using traditional statistical modeling approaches. Indeed, to date, deep learning have been used to generate photo-realistic cell images [3], learn functional representations of neural in-situ hybridization images [4], to predict novel drug targets [5, 6, 7, 8], and to model the hierarchical structure and function of the cell [9]. Recently, VAE methods have also been shown to learn biologically-relevant latent representations of tumors using genome-scale gene expression [10] and DNA methylation data11. In the application of DNA methylation, VAEs have been shown to learn latent representations of the breast tumor epigenome, and that this latent space contained biologically relevant information about intrinsic molecular tumor subtypes [11]. However, these applications are only beginning to emerge and interpreting the specific biology underlying learned latent dimensions of a VAE model remains a challenge.

Here, we train a VAE model on genome-wide DNA methylation data in breast tumors and extract sets of specific epigenomic features that contribute to the learned latent dimensions representing Estrogen Receptor (ER)-negative and ER-positive tumors. In breast cancer, distinct biology relating to tumor hormone status is a major factor in determining the treatment options available to patients, as ER-status is the main indicator of potential responses to endocrine therapy [12]. To demonstrate the feasibility of interpreting VAE models and extracting specific features of the input data, we show that the VAE model is capable of learning known biological and regulatory features representative of ER-status. These features, once learned, may have the potential to be use as methods for biological hypothesis generation and clinical risk stratification.

## Results

To investigate the feasibility of using variational autoencoder (VAE) models to extract biologically meaningful features from DNA methylation data, we trained a VAE model on the top 100,000 most variable CpGs across 1, 230 samples from three publicly available data sets, as defined by median absolute deviation (MAD) of methylation beta values (Table 1). The VAE model learned a 100-dimensional latent representation of the original 100,000-dimensional input data, representing each of the 100,000 original data inputs as a non-linear combination of the 100 intermediate dimensions. This 100D representation of the input data is refined by sampling from the 100 dimensions to attempt to reconstruct the original 100, 000 CpG methylation values. These 100 intermediate dimensions are referred to here as the latent dimensions of the model, and the trained VAE model is optimized to generate a lower-dimensional representation of the target DNA methylation data.

**Table 1.**
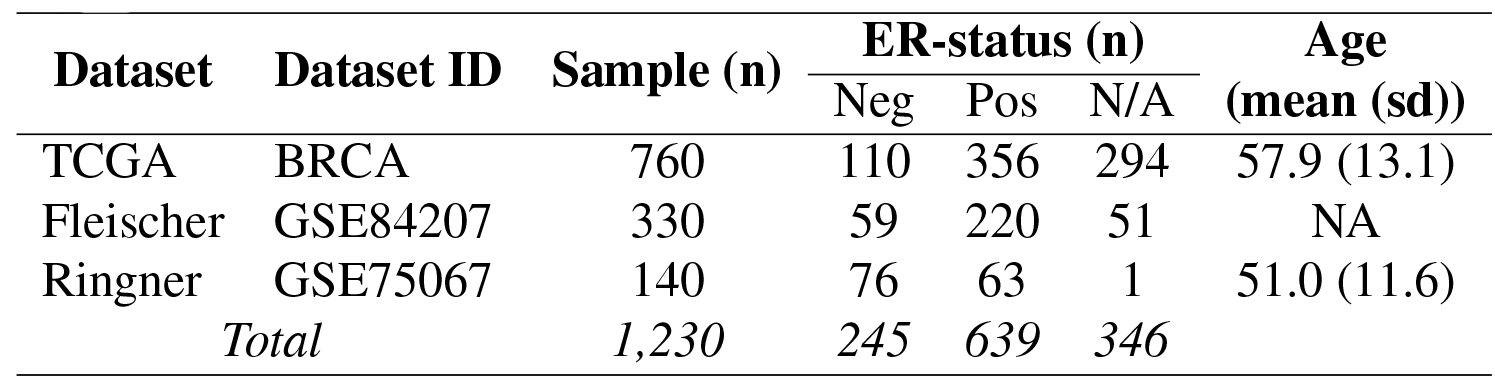
Breast tumor subject demographics, tumor characteristics, and DNA methylation data sources.

In breast cancer, distinct biology relating to tumor hormone status is a major factor in determining the treatment options available to patients, and estrogen receptor (ER)-status is the main indicator of potential responses to endocrine therapy [12]. To evaluate the efficacy of the VAE model to learn meaningful biology, we selected latent dimensions uniquely activated in either ER-negative or ER-positive breast tumors. These latent dimensions were further compared with the most highly activated dimensions across both tumor types. For dimensions representing all tumor types, the three dimensions with the most total activation were selected (dimensions VAE63, VAE37, VAE22) (Figure 1a, Supplemental Figure 1a). The three dimensions with the highest activation values in the respective tumor type (ER-/ER+) and a 75-percentile activation of zero in the contrasting tumor type were selected (ER-negative: dimensions VAE43, VAE35, and VAE24; ER-positive: dimensions VAE93, VAE91, and VAE47) (Figure 1b & 1c, Supplemental Figure 1b & 1c).

**Figure 1.**
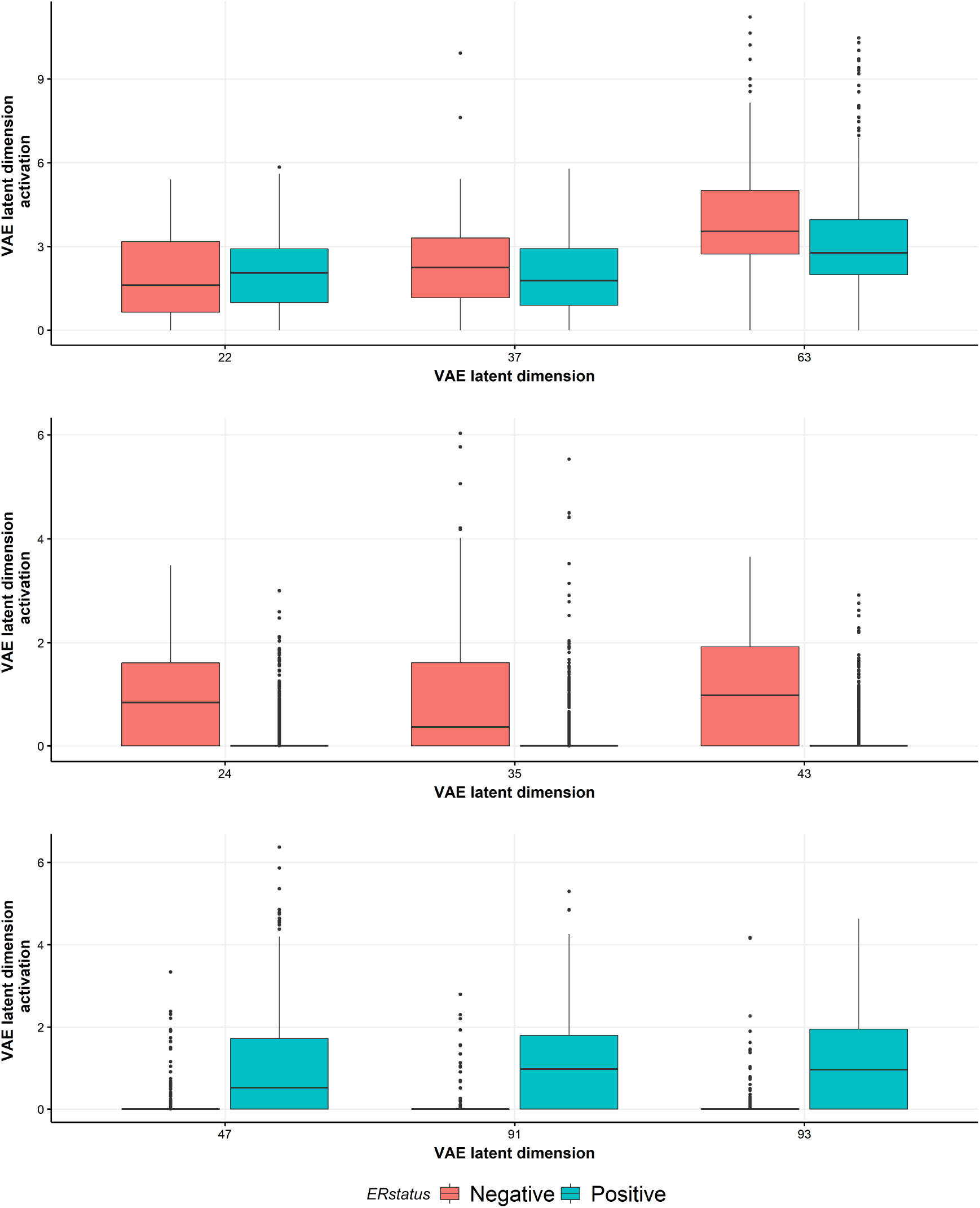
VAE latent dimensions with the highest activation values stratified by ER-status. The three latent dimensions representing each tumor class with the (a) highest overall activation values among all strata (dimensions VAE22, VAE37, an VAE63), and where (b) the VAE dimensions whose latent activation values was highest in ER-negative tumors and the 75th percentile of latent dimension activation value was zero in the ER-positive tumors (dimensions VAE24, VAE35, and VAE43), and (c) where the VAE dimensions whose latent activation values were highest in ER-positive group and the 75^*th*^ percentile of latent dimension activation value was zero in the ER-negative tumors (dimensions VAE47, VAE91 and VAE93).

After identifying VAE latent dimensions with differential activation values between ER-negative and ER-positive tumors, we sought to identify CpG sites with the strongest association of DNA methylation state activation values for each latent dimension. Pairwise Spearman correlation tests between the methylation state of each CpG and the activation values of each of the nine selected latent dimensions were performed. Then, to select a subset of CpGs for further investigation we used a threshold correlation value for each latent dimension based on the inflection point at which the correlations began to rapidly decrease. The final set of CpGs for subsequent analyses for each of the nine latent dimensions are shown in Table 2, and Supplemental Figure 2. We imposed a minimum of 50 CpGs with high latent dimension activation value correlation for subsequent genomic context analyses (Supplemental Figure 2b & 2c).

**Table 2.**
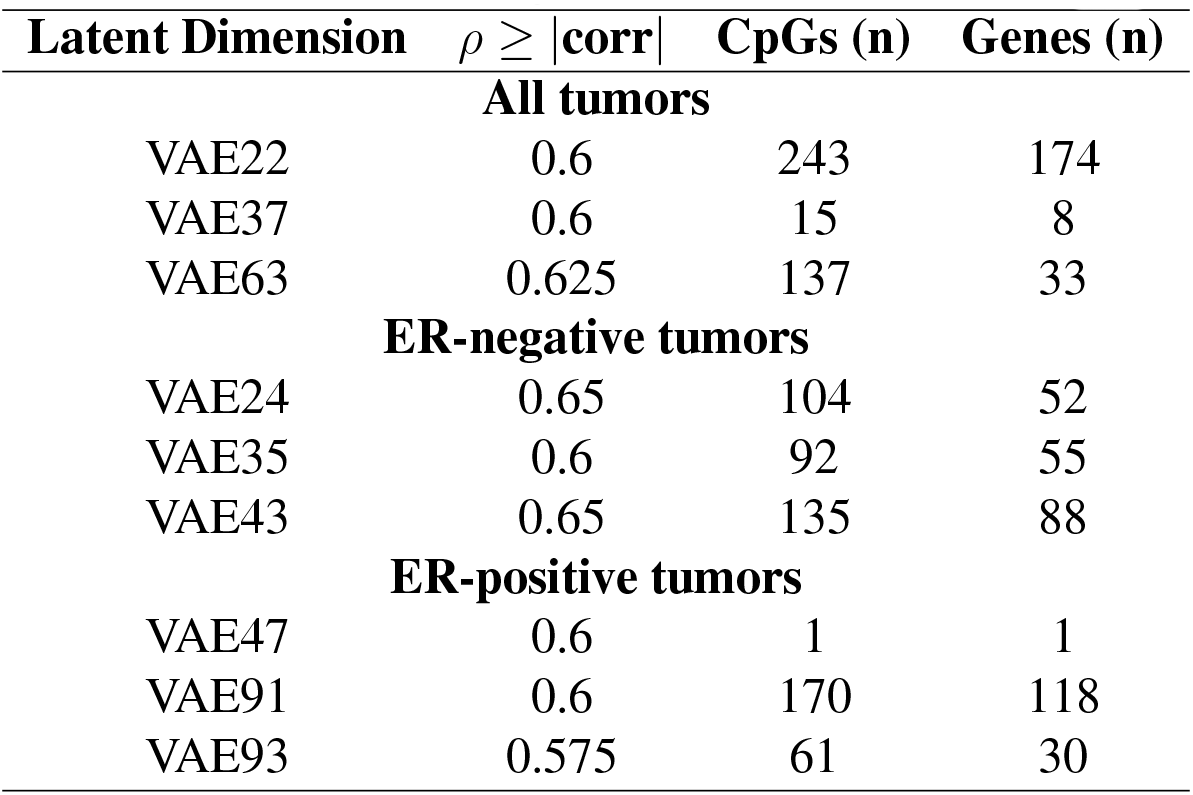
Correlation of CpG methylation with latent dimension activation.

From the remaining seven latent dimensions (All tumors: VAE22, VAE63; ER-negative: VAE24, VAE35, VAE43; ER-positive: VAE91, VAE93) we investigated where in the genome the identified set of CpGs were located. In those dimensions associated with all tumors (VAE22, VAE63) and those associated with ER-negative tumors (VAE24, VAE35, VAE43) and ER-positive tumors (VAE91, VAE93), the identified loci were genome-wide and showed no obvious genomic distribution (Figure 2, Supplemental Figure 3, Supplemental Data 1-7).

**Figure 2.**
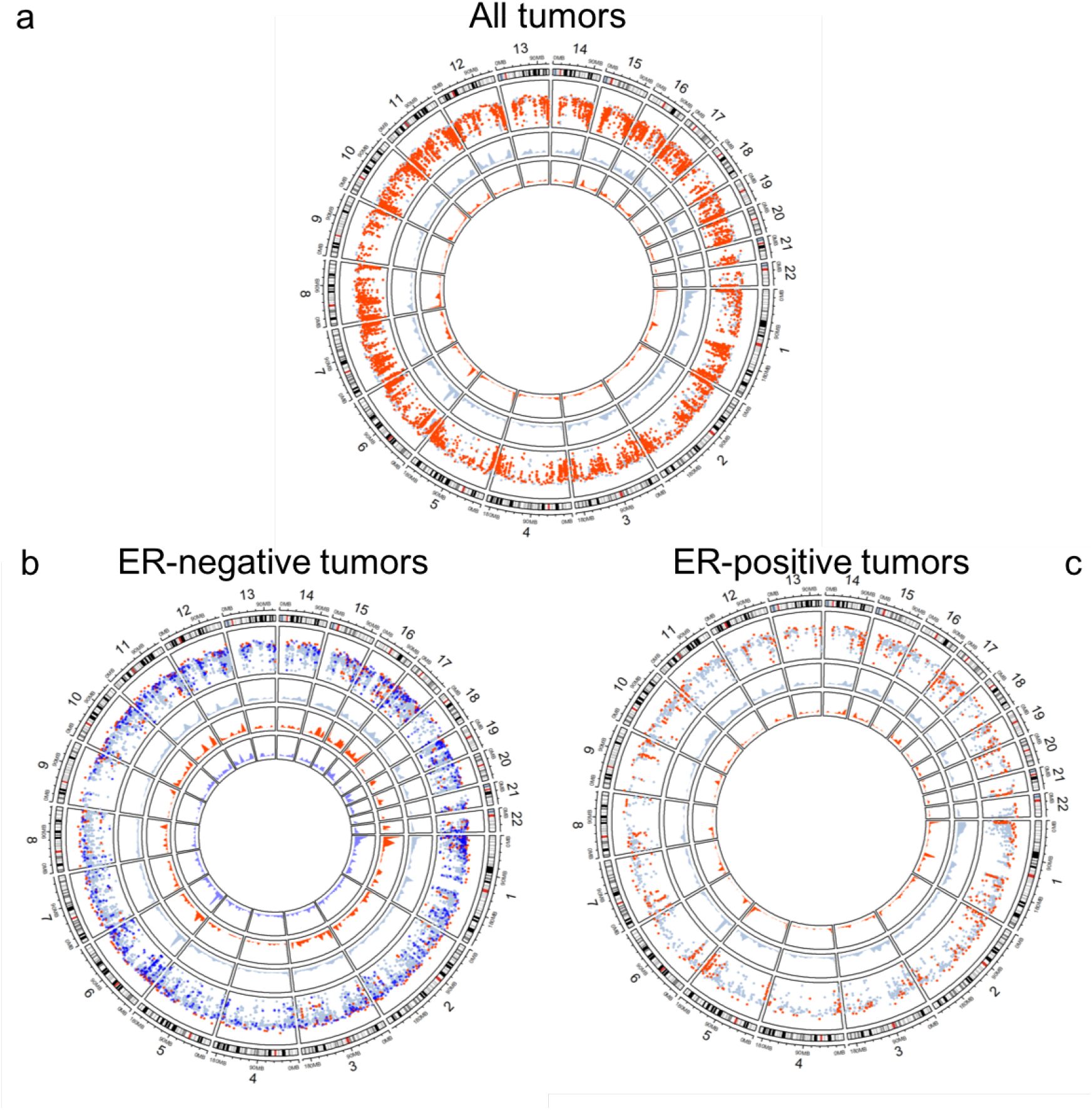
Genomic context of CpGs most correlated with VAE latent dimensions. The outer ring displays chromosome location, the next inner ring shows the individual CpGs for all three dimensions, and the inner most rings display the relative CpG densities. (a) All tumors (VAE22 light blue, VAE63 red), (b) ER-negative tumors (VAE24 light blue, VAE35 red, VAE43 dark blue), and (c) ER-positive tumors (VAE91 light blue, VAE93 red).

To test the hypothesis that VAE-correlated CpG loci are enriched among pathways related to ER-status in breast cancer, we conducted gene ontology (GO) and gene set enrichment (GSEA) analyses. Due to a lack of CpG sample size, and by extension gene annotations (Supplemental data 1-7), most latent dimension CpGs sets did not reach an FDR *<* 0.05 significance threshold when conducting GO term analyses. However, the CpG set associated with VAE43 (ER-negative tumors) identified three significant immune related GO terms (regulation of CD8-positive alpha-beta T cell activation, GO:2001185, FDR = 0.002; adaptive immune response, GO:0002250, FDR = 0.002; CD8-positive, alpha-beta T cell activation, GO:0036037, FDR = 0.04) (Supplemental Data 8-14). In the GSEA analysis of the CpGs sets from all tumor related dimensions (VAE22, VAE63), our analysis identified twenty FDR *<* 0.05 significant pathways related to VAE22, with the most significant pathway being for luminal versus mesenchymal breast cancer (FDR = 2.76*e* 04, Table 3) and no pathways with FDR *<* 0.05 for VAE63 (Table 3, Supplemental Data 15-16). In the GSEA analysis of the CpGs sets from ER-negative related dimensions (VAE24, VAE35, VAE43), our analysis identified forty-six significant pathways related to VAE24 (FDR P *<* 0.05), with the most significant pathway being target genes for STAT3 in CSF3 signaling (FDR = 0.008, Table 3). There were nine significant pathways (FDR *<* 0.05) related to VAE35, with the most significant pathway being for genes upregulated when E2F3 is knocked down (FDR = 0.011, Table 3). Lastly, there were over fifty significant pathways (FDR *<* 0.05), related to VAE43, with the most significant pathway being for genes upregulated with IFN-*β* treatment (FDR = 2.86*e* 05, Table 3, Supplemental Data 17-19). In the GSEA analysis of the CpGs sets from ER-positive related dimensions (VAE91, VAE93), our analysis identified ten significant pathways related to VAE91 (FDR *<* 0.05), with the most significant pathway being for genes shown to have copy number loss in primary neuroblastomas (FDR = 6.43*e* 05, Table 3) and one significant pathway related to VAE93 (FDR *<* 0.05), with the most significant pathway being for linoleic acid metabolism genes (FDR = 0.027, Table 3, Supplemental Data 20-21).

**Table 3.**
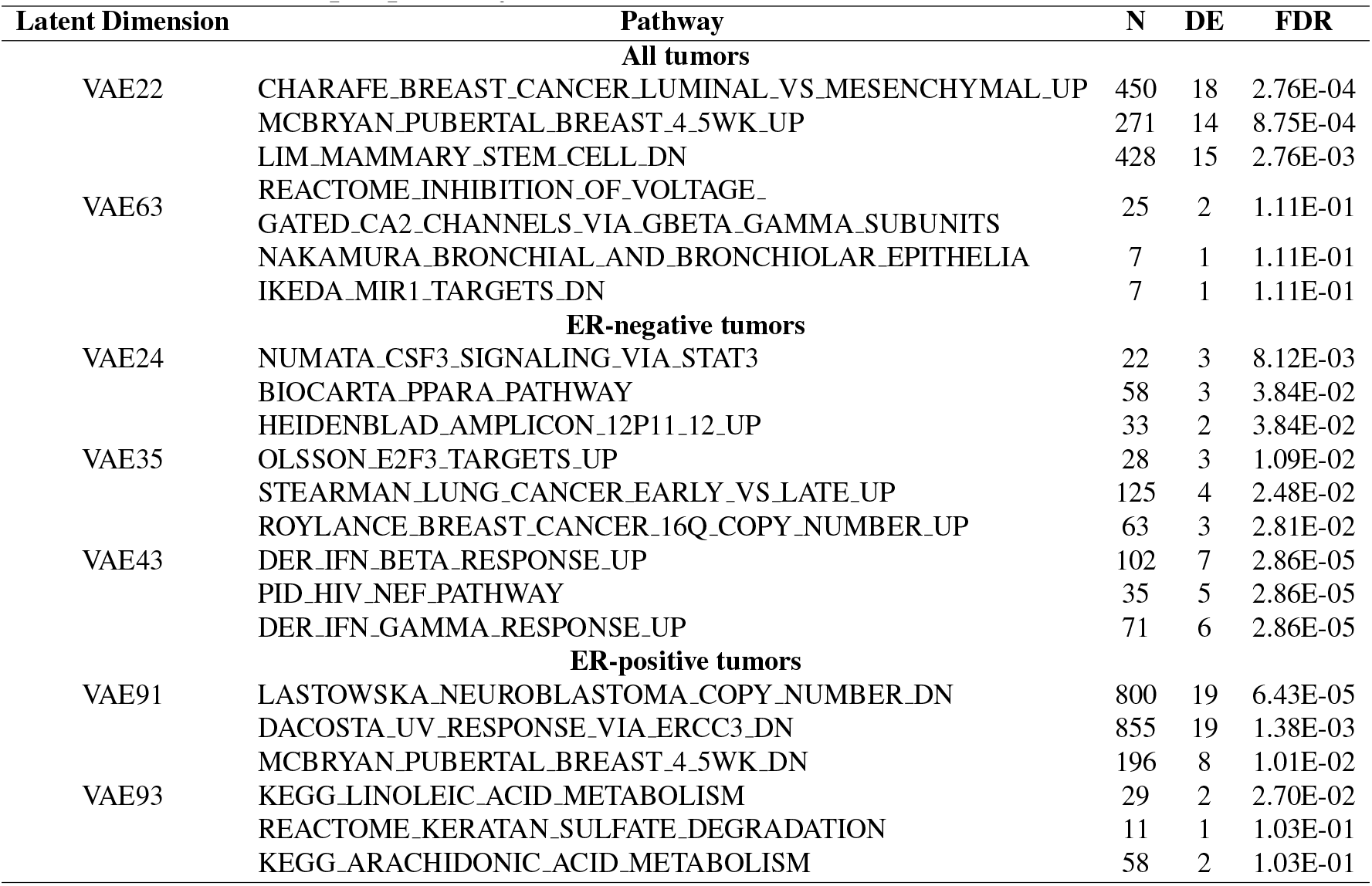
Top 3 pathways associated with each VAE latent dimension.

After identifying the set of all CpGs with a correlation between methylation and latent dimension activation above the threshold, we identified the individual CpGs most correlated with each dimension (VAE24:cg24920933, VAE35:cg04218548, VAE43:cg03032552, VAE91:cg00238899, VAE93:cg19103219, VAE22:cg24486037, VAE63:cg03812562) (Figure 3). Of the seven CpGs identified as most corelated with each latent dimension, only three were annotated to genes (VAE35 – Chr1:*CAMTA1*, VAE93 – Chr10:*MXI1*, VAE22 – Chr1:*SELENBP1*).

**Figure 3.**
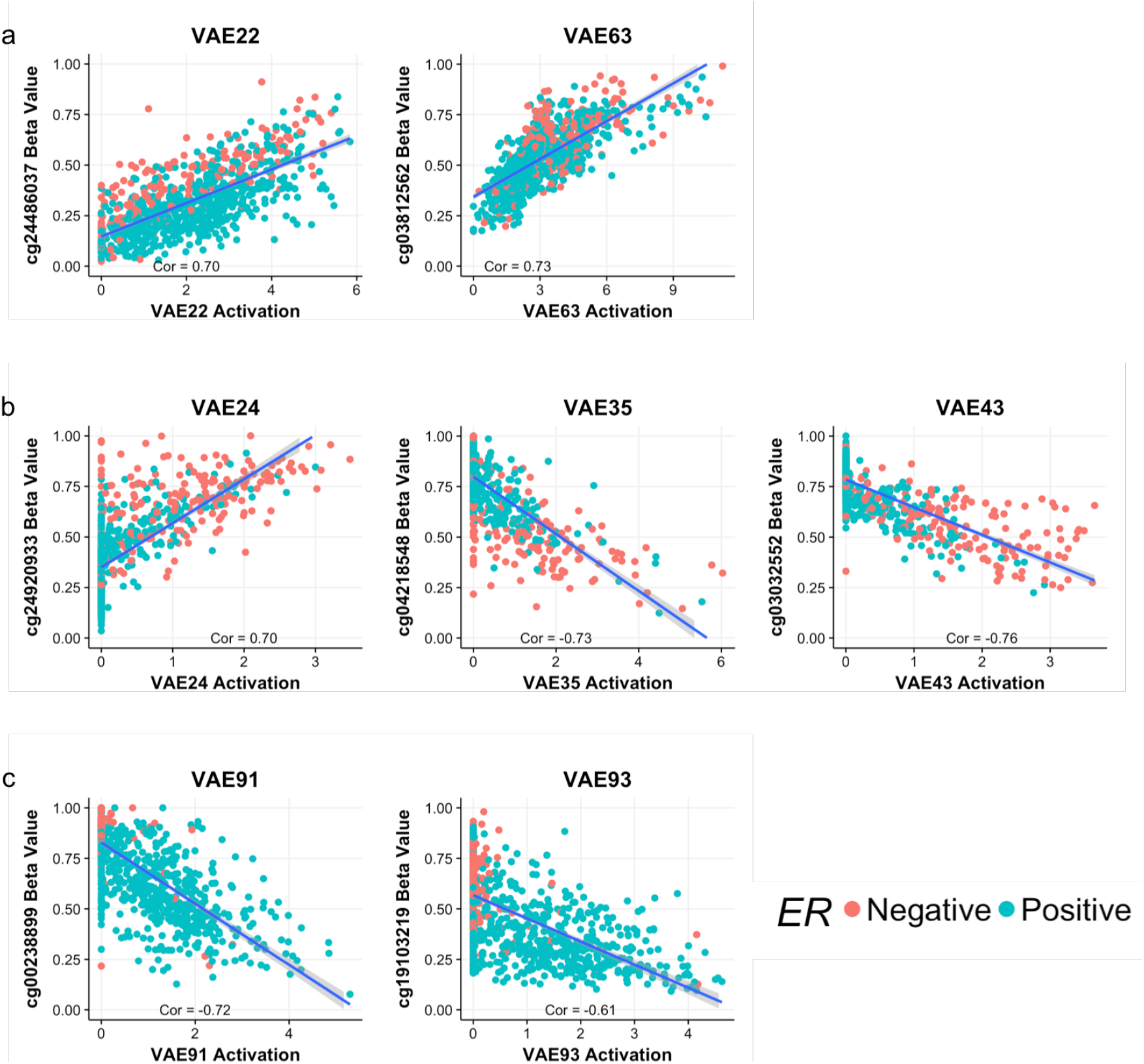
Correlation of CpG methylation with VAE latent dimension activation values for each selected latent dimension at the CpG with the greatest absolute correlation value within the respective analysis group (all tumors, ER-negative tumors, ER-positive tumors). Red points represent ER-negative samples and blue points represent ER-positive samples. Row (a) represents all-tumor associated latent dimensions, row (b) represents ER-negative associated latent dimensions and row (c) represents ER-positive associated latent dimensions.

With five out of the seven CpGs most correlated being unannotated to genes, we investigated the proportion of all identified CpGs that were annotated to genes. We found that the proportion of CpGs associated with each latent dimension varied from 0.28 annotated to 0.78 annotated (VAE22 = 0.78, VAE63 = 0.28, VAE24 = 0.51, VAE35 = 0.70, VAE43 = 0.76, VAE91 = 0.75, VAE93 = 0.58) (Table 4).

**Table 4.** CpG with highest correlation between methylation value and latent dimension activation value, and the proportion of all CpGs within each set with known gene annotations.

| Latent Dimension | CpG n (%) with Gene Annotations | CpG n (%) without Gene Annotations |
| --- | --- | --- |
| **All tumors** |  |  |
| VAE22 | 190 (78) | 53 (22) |
| VAE63 | 39 (28) | 98 (72) |
| **ER-negative tumors** |  |  |
| VAE24 | 53 (51) | 51 (49) |
| VAE35 | 64 (70) | 28 (30) |
| VAE43 | 103 (76) | 32 (24) |
| **ER-positive tumors** |  |  |
| VAE91 | 128 (75) | 42 (25) |
| VAE93 | 36 (59) | 25 (41) |

Given that a major functional determinant of DNA methylation is CpG island context (ocean, shore, shelf, or sea), we tested for enrichment of VAE correlated loci with CpG island context. Six of the seven analyzed VAEs were strongly significantly depleted for CpG island annotations and were significantly enriched for CpG open sea annotations (VAE22, VAE24, VAE35, VAE43, VAE91, VAE93) (Figure 4, Supplemental Data 22), suggesting a concentration of CpGs among non-coding, CpG sparse regions.

**Figure 4.**
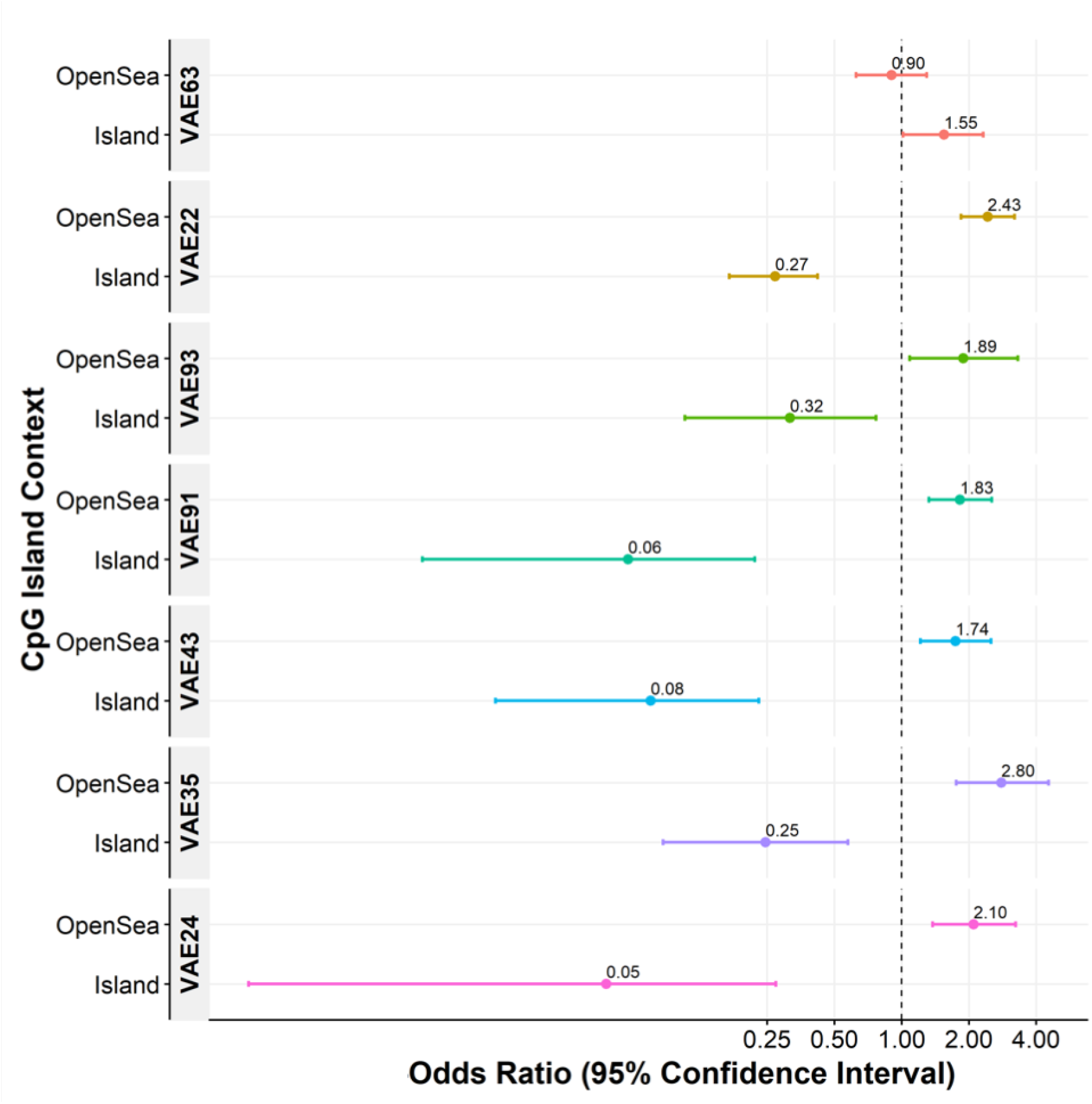
Forest plot showing the genomic context enrichment of latent dimension-related CpG sets. CpGs related with latent dimension activation values are consistently depleted for CpG islands and enriched for open sea regions. Enrichment tests were conducted with Cochran-Mantel-Haenszel tests, controlling for Illumina probe type.

Intergenic DNA, and effective functional annotation of these regions among distinct cell-types remains challenging. To determine if VAE correlated CpG loci are associated with specific DNA regulatory regions, we conducted tests using Cochran-Mantel-Haenszel, controlling for Illumina probe type, to test for enrichment of non-cell-type-specific enhancer elements and DNase 1 hypersensitivity sites (DHS), as annotated by the ENCODE consortium [13, 14]. Four out of seven latent dimensions were significantly enriched for enhancer annotations (VAE22, VAE91, VAE35, VAE24), and one latent dimension was significantly depleted for enhancer annotations (VAE63) (Figure 5). One VAE latent dimension was significantly enriched (VAE22) and three latent dimensions were significantly depleted (VAE63, VAE93, VAE24) for DNase 1 hypersensitivity sites (Figure 5, Supplemental Data 22).

**Figure 5.**
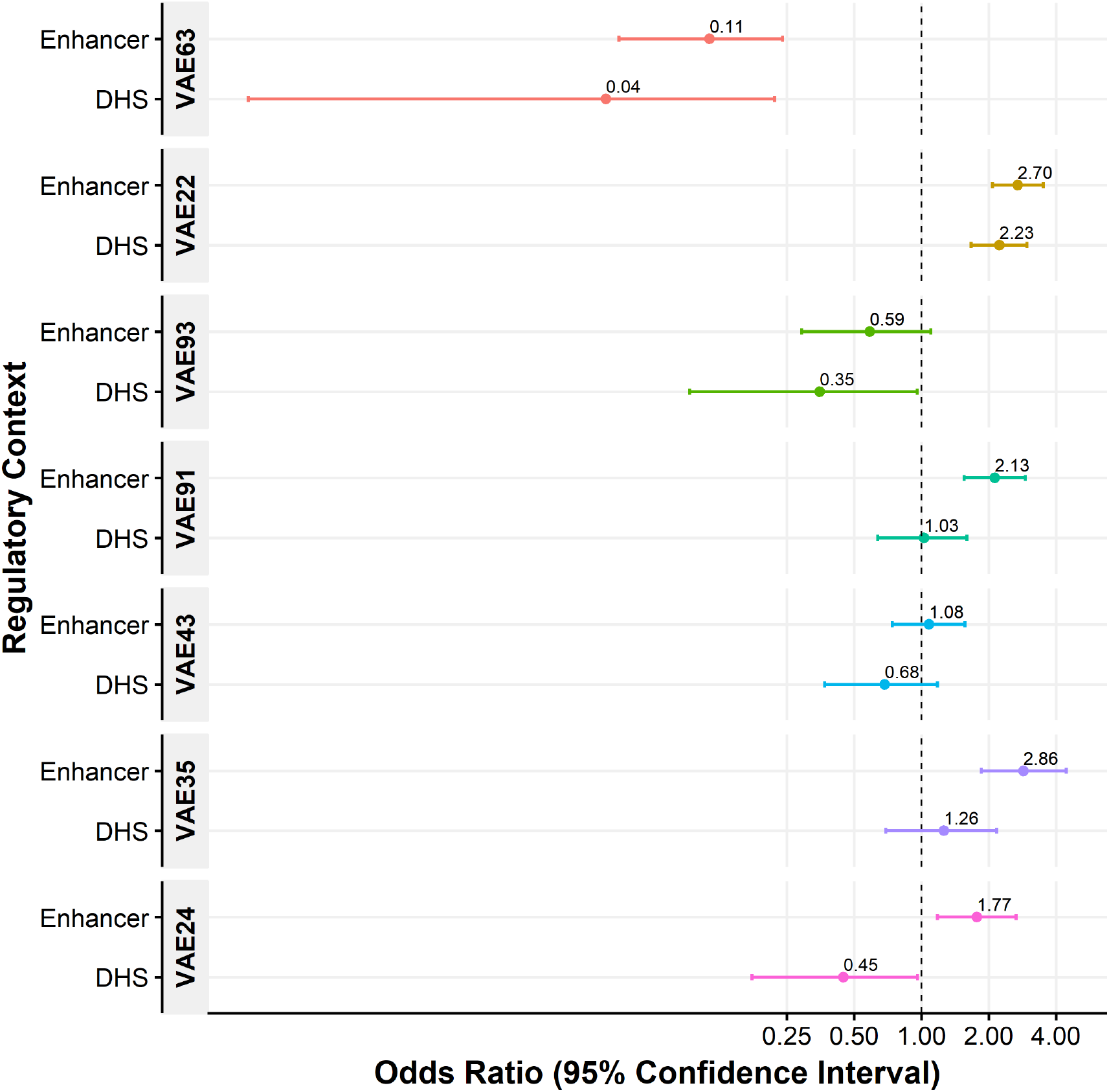
Enrichment analysis of latent-dimension-related CpG sets for (a) enhancer annotations and (b) DNase 1 hypersensitivity site annotations. Enrichment tests were conducted with Cochran-Mantel-Haenszel tests, controlling for Illumina probe type.

With evidence of cell-type agnostic enhancer enrichment, we sought to assess breast tissue specific regulatory enrichment. Using ENCODE consortium ChIP-seq data from breast myoepithelial cells (Br myo) and human mammary epithelial cells (HMEC), we conducted enrichment analyses for histone modification marks associated with transcriptional repression (Br myo: H3K4me1, H3K4me3, H3K36me3; HMEC: H3K9me3, H3K20me1, H3K27me3) and histone modification marks associated with transcriptional activation (Br myo: H3K9ac, H3K27me3; HMEC: H3K4me1, H3K4me2, H3K4me3, H3K9ac, H3K27ac, H3K36me3, H3K79me2). Among the latent dimensions, those associated with all tumors were split between enrichment (VAE22) and depletion (VAE63) for transcriptional repressive marks. A similar pattern was observed among the latent dimensions representing ER-negative tumors, with dimensions representing depletion (VAE24), enrichment (VAE35), and mixed enrichment and depletion (VAE43) for transcriptionally repressive marks. The latent dimensions representing ER-positive tumors were mixed with both enrichment and depletion of transcriptionally repressive marks (VAE91, VAE93). In general, however, five latent dimensions were depleted (VAE24, VAE43, VAE91, VAE93, VAE63), and only two were enriched for transcriptional activating marks (VAE22, VAE35) (Figure 6, Supplemental Data 22).

**Figure 6.**
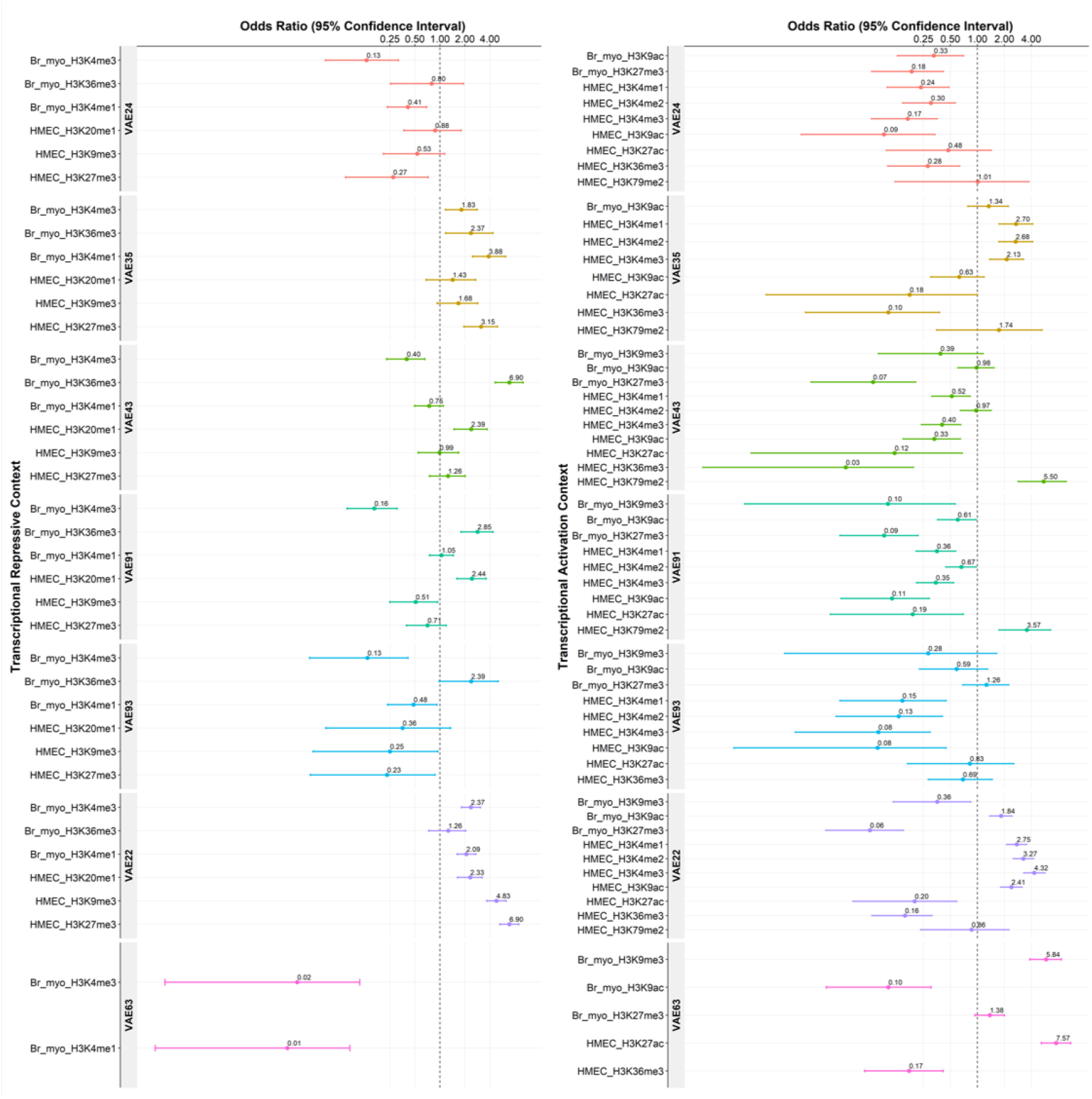
Enrichment analysis of latent dimension-related CpG sets for histone marks related to (a) transcriptional repression and (b) transcriptional activation. Enrichment tests were conducted with Cochran-Mantel-Haenszel tests, controlling for Illumina probe type.

To investigate if there is redundancy with common approaches for dimensionality reduction and feature selection, we compared the VAE model CpG set to loci selected using LASSO regression, comparing ER-negative to ER-positive tumors. With the VAE model, we identified 562 CpGs (560 unique loci) associated with the five latent VAE dimensions related to ER-negative (VAE24, VAE35, VAE43) and ER-positive tumors (VAE91, VAE93). The LASSO regression identified 146 CpGs as the set of CpGs most discriminative between ER-negative and ER-positive tumor samples. The VAE and LASSO analyses identified 3 of the same CpGs, representing 0.5% of the VAE identified CpGs and 2.1% of the LASSO identified CpGs. The lack of overlap between the two CpG sets suggest that the VAE model is identifying non-redundant information about CpG DNA methylation patterns in breast cancer.

## Discussion

Deep learning is a promising approach for the study of complex biological systems, including the epigenome, but issues remain regarding our ability to delineate the learned biology from these models. There are open questions about what the intermediate, hidden layers within deep learning models are doing during training, but the lack of clarity is often acceptable in supervised models when performing successful predictions because the models rely on *a priori* data annotations. Whereas here, with unsupervised methods, we can build models but then need to interrogate the latent dimension to make interpretations. We trained a variational autoencoder (VAE)-based deep learning model on genome-wide breast cancer DNA methylation data to test the relation of latent dimension activation values with tumor phenotypes. In addition, to facilitate model interpretation, we outline approaches for interrogating the biology most related with latent dimension activation values. Through this framework we identified enrichment for CpGs most related with latent dimension activation values to CpG sparse regions of the genome, and specifically to enhancer elements.

VAEs pose many distinct advantages over traditional analytical methods when exploring the complex nature of the epigenome. First, the models are unsupervised and thus are entirely data driven and do not rely on existing genomic annotations. The reliance on existing annotations limits our ability to uncover novel biology and has been shown to push researchers towards known biology over that with the strongest molecular data [15]. As a data drive model, deep learning VAE models repeatedly adjust a weighted combination of input features until the model identifies the best possible reconstruction of the input data. This can be thought of as a high dimensional interaction space in which the latent dimensions capture genome-wide sets of CpGs that are similarly associated with ER-status, histone modifications, and gene sets.

Our initial effort to use deep learning with breast tumor DNA methylation data to recapitulate intrinsic breast tumor molecular subtypes established the feasibility and promise of unsupervised deep learning approaches for molecular data sets [11]. Other work has begun to use unsupervised deep learning to investigate shared transcriptomic signatures in a pan-cancer setting [10], and to identify a potential response to drug treatments and drug-drug interactions [5, 6, 7, 8]. There is potential, then, to use latent representations of tumors in this way to predict a patient’s response to therapies, or to stratify patients by latent risk profile. The learned latent representation of the breast tumor methylome can also be used for inference on new data samples. ER-status, for example, must be measured but often this is a missing piece of data, such as in the 346 samples with missing ER-status in this study. Because the VAE is data driven and unsupervised, the training process condenses all information about a tumor into a lower dimensional space and can thus be used for inference of novel information.

The most striking model results of our analysis is the wide-spread enrichment and/or depletion of histone modifications and enhancers across the identified latent dimensions, which is consistent with existing literature. Our VAE model identified modification HMEC-H3K9me3 significantly depleted in both latent dimensions associated with ER-positive tumors (VAE91, VAE93) and enriched in one latent dimension related to all tumors. Similarly, the model identified Br myo-H3K9me3 as significantly depleted in one latent dimension related to ER-positive tumors (VAE91) and one related to all tumors (VAE22) as well as enriched in one latent dimension related to all tumors (VAE63). This is consistent with studies showing estrogen-induced rat breast carcinogenesis is characterized by rapid and sustained loss of H3K9me3 in breast tissue [16]. Similarly, the wide-spread role H3K4 and H3K27 histone marks play in cancer has been previously discussed [17], and our VAE model identifies latent dimensions associated with both significant depletion and enrichment for CpG loci associated with H3K4me1 (HMEC, Br myo), H3K4me2 (HMEC), H3K4me3 (HMEC, Br myo), H3K27me3 (HMEC, Br myo), and H3K27ac (HMEC).

These findings suggest the most representative epigenomic features in the training process lie within DNA regulatory regions. Given that epigenetic regulation is a critical determinant of cellular function in adult tissues [18, 19], this observation supports the hypothesis that VAEs identify a biologically meaningful latent space from DNA methylation data. Further supporting these findings, ubiquitous enrichment of the learned latent dimensions amongst open sea regions supports the conclusion that that intergenic DNA regulatory regions play a key role in the model training process. Historically, DNA methylation analyses focus on CpG dense promoter regions of the genome, but the results extracted from the learned VAE model suggest that additional benefit may be gained by expanding the analysis space to include CpG sparse regions of the genome in future analyses. These results are consistent with recent work identifying partially methylated domains, or ‘solo-WCGW’, in CpG-sparse regions of the genome associated with cancer [20]. Additionally, as these models can be thought of as latent high dimensional interaction spaces, we may begin to employ such techniques to study the 3-dimensional nature of the genome, investigating both local and long-range chromosomal interactions as well as local regulatory domains, such as tandem associating domains (TADs). There may also be opportunities to use analytical approaches such as this in supervised leaning models to investigate the nature of the learned biology. It is likely that deep learning models will become more accepted in the clinical setting due to lower error rates in predictions. Along with this adoption, we should continually try to understand and intervene when prediction errors are made so we might improve on the input data, or how its measured, in order to further reduce prediction error.

When investigating the genomic features associated with each of the seven identified latent dimensions, it became clear that many of the dimensions were learning complementary biology. Among the three latent dimensions activated in ER-negative tumors, VAE24 was most associated with a CpG set that was enriched for open sea contexts, depleted across HMEC-related histone modifications, and was most associated with the gene set related to CSF-3 signaling. CSF is immune related and has been shown to regulate macrophage phenotypes and is associated with poor survival in patients with triple negative breast cancer (ER-negative) [21]. In contrast to VAE24 depletions across HMEC histone modifications, VAE35 is enriched across the HMEC histone modifications while still maintaining an enrichment for open sea contexts. VAE35 is also associated with the upregulation of E2F target genes which is a gene set known to be an important player in breast cancer tumor development and progression [22]. E2F3 also plays a critical role in tumor macrophages and gains in E2F3 are more common in ER-negative breast cancer than in ER-positive breast cancer [23]. The final ER-negative related latent dimension, VAE43, was found to be enriched for open sea contexts and was significantly associated with GO terms related to immune function. This analysis indicates that the CpGs most associated with VAE43, and therefore ER-negative tumors, are significantly associated with immune regulation. In addition, VAE43 was found to be significantly associated with the gene set related to interferon beta response, which has been shown to be a promising target for triple negative breast tumors [24].

Within the identified latent dimensions related to ER-positive tumors, the VAE91 CpG set was enriched for open sea context and for enhancers and the GSE analysis identified VAE91 as significantly associated with the set focused around neuroblastoma copy number variation. Copy number variation has been shown to play a major role in breast cancer [23] and are associated with risk and prognosis [25]. The VAE93 dimension was also enriched for open sea context, but in contrast to VAE91 enhancer enrichment, VAE93 is significantly depleted of enhancers. VAE93 was also associated with linoleic acid metabolism, which has been shown to have growth-inhibitory and proapoptotic effects on ER-positive tumors.

The identified latent dimensions where complementary learned biology is the most apparent, were the two identified with the highest activation values across breast tumor subtypes. Latent dimensions VAE22 was enriched for open sea context, enhancers, and DNase Hypersensitivity Site 1. VAE22 also contained the largest proportion of identified CpGs across all identified latent dimensions, and similarly contained the greatest proportion of CpGs with known annotations to genes. VAE22 also was strongly associated with the gene set related to the upregulation of luminal breast cancer related genes relative to mesenchymal breast cancer related genes. In contrast, VAE63 was depleted of enhancers, DNase Hypersensitivity Site 1, and was the only one of the seven identified latent dimensions that was not enriched for open sea context and has a substantial portion of the CpGs located in CpG Islands. At the same time, 72% of the identified CpGs associated with VAE63 do not have a gene annotation. This indicates that while there are ample CpGs located in CpG islands, many of these islands are not located within a known gene annotation. This lack of specific gene annotation is also the reason that VAE63 was the only identified latent dimension that was not significantly associated with any gene set through the GSEA analysis. Overall, both VAE22 and VAE63 are associated with all tumor types, but they represent two latent dimensions capturing different sets of information, with VAE22 capturing gene information and VAE63 capturing intergenic information.

Features from VAEs have been used for a number of applications in recent years, yet there is still a lack of interpretability inherent in deep learning models due primarily to the non-linear nature of the model architectures. Efforts are ongoing to make deep learning models more interpretable, more accessible, and more useful to biologists [1]. Here we present a framework for training a deep learning model on genome-wide DNA methylation data and demonstrate how biologically meaningful insights can be extracted from such a model.

## Methods

### Data

Three genome-wide breast cancer DNA methylation data sets were used during model training. We downloaded all Illumina HumanMethylation450 (450K) DNA methylation level 1 sample intensity data files for breast invasive carcinoma (*n* = 760) from The Cancer Genome Atlas (TCGA) data access portal [26]. We processed the data files with the R package minfi [27] using the Funnorm normalization method for the full dataset. For quality controls, we filtered CpGs with a detection P-value *>* 1.0*e* – 05 in more than 25% of the samples, CpGs with high frequency SNP(s) (*>* 5% minor allele frequency) in the probe, probes previously described to be potentially cross-hybridizing, and sex-specific probes. From an original set of 485, 512 measured CpG sites on the 450K array, our filtering steps removed 2, 932 probes exceeding the detection P-value, and 93, 801 probes that were SNP-associated, cross-hybridizing, or sex-specific resulting in a final analytic set of 388, 779 CpGs (Table 1).

We also downloaded two publicly available breast cancer data sets from the Gene Expression Omnibus (GEO) measured on the Illumina 450K platform, one provided by Fleischer *et al.* (*n* = 330, GSE84207) [28], in which data underwent standard pre-processing and normalization with probe filtering, color bias correction, background subtraction and subset quantile normalization [29]. We also included data provided by Ringner *et al.* (*n* = 140, GSE75067) [30], in which methylation beta values were obtained using Illumina GenomeStudio, and samples with detection p values greater than 0.05 or with a fewer than three beads per channel were treated as missing data. The normalized methylation values were used as deposited in GEO, resulting in a final data set of 1, 230 samples (Table 1).

To reduce the number of input dimensions for our model, the top 100, 000 variable CpGs, measured by median absolute deviation (MAD) [31] of methylation value across all three data sets, were used for model training [10].

### Variational Autoencoder Model

We extend the *Tybalt* VAE model to learn a latent methylome from genome-wide DNA methylation data. *Tybalt* was developed by Way *et al.* to learn latent pan-cancer transcriptomes [10]. The model consists of an Adam optimizer [32], rectified linear units [33], and batch normalization in the encoding stage, and sigmoid activations in the decoding stage. The model is built in Keras (Version 2.0.6) [34] with a Tensorflow backend (version 1.0.1) [35, 36]. We trained the model using optimal parameters identified by Way *et al.* [10] with the following values: batch size = 50, learning rate = 0.0005, k= 1, epochs = 50, test/validation = 90/10.

The original model was designed for 5, 000 input genes encoded to 100 latent dimensions. We adapted the model to take in and reconstruct 100, 000 CpG methylation values, proportion of alleles methylated at a specific site, with 100 latent dimensions. The 100, 000 CpGs were selected based on highest variability by median absolute deviation (MAD) [31] of methylation values. The output of the VAE model results in a 100-dimensional latent feature vector for each study sample, where the specific value of each of the 100 dimensions is referred to as the latent dimension’s “activation” value.

### Latent Dimension Analyses

Using the estrogen receptor (ER) status of tumors, we selected the three latent dimensions most associated with all tumors, as well as three representing each of ER-negative and ER-positive tumors individually, for subsequent analyses. The three dimensions with the greatest median total activation across all samples were selected as most associated across all tumors. Additionally, because these final three dimensions were selected to represent all ER-statuses, we included those tumors with unknown ER-status. Three latent dimensions most associated with ER-negative and ER-positive tumors each were selected by choosing the latent dimensions with the greatest total activation across all samples in the target tumor type (ER-positive or ER-negative), and an activation 75th-percentile of zero in the comparison tumor type. In other words, each set of three latent dimensions were selected to ensure that they represent the most activation possible in the target tumor type, while 75% of the samples in the alternative tumor type had an activation value of zero.

The relationships of input features with hidden layers within a deep learning model are often non-linear and challenging to interpret. To identify specific CpGs meaningfully-associated with selected latent dimensions, pairwise Spearman correlations were calculated between each of the 100, 000 input CpG methylation values and the activation states of the nine selected latent dimensions. Spearman correlation is a non-parametric method to measure the strength and direction of association between two ranked variables. In this case, the absolute value of the correlation was assessed in a ranked order relative to the activation values of the latent dimensions and a threshold was set for each latent dimension based on the inflection point at which the correlations began to rapidly decrease. CpGs with correlations above the respective threshold were kept for subsequent analyses.

After identifying the set of CpGs most correlated with the latent dimensions, we investigated where the identified CpGs were distributed throughout the genome, both combined by ER-status and individually across chromosomes. We also conducted gene ontology (GO) term and gene set enrichment analyses (GSEA) on sets of CpGs most correlated with latent dimensions using the R package missMethyl [37]. Sets of CpGs most correlated with latent dimensions were then tested for enrichment with gene regulatory features including Enhancers and DNase Hypersensitive Sites 1 based on the Illumina HumanMethylation450 annotation file (Enhancer & DHS) using Cochran-Mantel-Haenszel tests, controlling for Illumina probe type (Type I & II). The enhancer annotations from the Illumina annotation file are provided by the ENCODE consortium [13, 14] and are not tissue specific. To investigate breast tissue specific regulatory regions, we conducted further enrichment tests among regulatory region DNA using Cochran-Mantel-Haenszel tests, controlling for Illumina probe type, for histone marks associated with transcriptional repression using ChIP-seq data from breast myoepithelial cells (Br myo) and human mammary epithelial cells (HMEC) (Br myo: H3K4me1, H3K4me3, H3K36me3; HMEC: H3K9me3, H3K20me1, H3K27me3) as well as histone marks associated with transcriptional activation (Br myo: H3K9ac, H3K27me3; HMEC: H3K4me1, H3K4me2, H3K4me3, H3K9ac, H3K27ac, H3K36me3, H3K79me2) downloaded from the ENCODE data repository [13, 14].

To compare the CpGs identified using the VAE model to standard feature selection methods, we also conducted LASSO regression to identify the CpG features most discriminative between ER-negative and ER-positive tumors.

## Supplemental Data

Supplemental data from this analysis can be found at the following location: https://doi.org/10.5281/zenodo.1442905

## Code Availability

All code used for this analysis can be found on GitHub at the following location: https://github.com/Christensen-Lab-Dartmouth/VAE_methylation.

## Acknowledgements

Research reported in this publication was supported by the Office of the U.S. Director of the National Institutes of Health under award number T32LM012204 to AJT, grants R01DE022772 and R01CA216265 to BCC, and by a Burroughs Wellcome Fund fellowship to CAB under award number #1014106. This research was also supported by The Dartmouth Clinical and Translational Science Institute, under award number UL1TR001086 from the National Center for Advancing Translational Sciences (NCATS) of the National Institutes of Health (NIH) and by an award from the Dartmouth College Neukom Institute for Computational Science. The content is solely the responsibility of the author(s) and does not necessarily represent the official views of the NIH.

